# Stratified microbial communities in Australia’s only anchialine cave are taxonomically novel and drive chemotrophic energy production via coupled nitrogen-sulphur cycling

**DOI:** 10.1101/2023.04.03.535450

**Authors:** Timothy M. Ghaly, Amaranta Focardi, Liam D. H. Elbourne, Brodie Sutcliffe, William Humphreys, Ian T. Paulsen, Sasha G. Tetu

## Abstract

**Background:** Anchialine environments, in which oceanic water mixes with freshwater in coastal aquifers, are characterised by stratified water columns with complex physicochemical profiles. These environments, also known as subterranean estuaries, support an abundance of endemic macro and microorganisms. There is now growing interest in characterising the metabolisms of anchialine microbial communities, which is essential for understanding how complex ecosystems are supported in extreme environments, and assessing their vulnerability to environmental change. However, the diversity of metabolic strategies that are utilised in anchialine ecosystems remains poorly understood.

**Results:** Here, we employ shotgun metagenomics to elucidate the key microorganisms and their dominant metabolisms along a physicochemical profile in Bundera Sinkhole, the only known continental subterranean estuary in the Southern Hemisphere. Genome-resolved metagenomics suggests that the communities are largely represented by novel taxonomic lineages, with 75% of metagenome-assembled genomes assigned to entirely new or uncharacterised families. These diverse and novel taxa displayed depth-dependent metabolisms, reflecting distinct phases along dissolved oxygen and salinity gradients. In particular, the communities appear to drive nutrient feedback loops involving nitrification, nitrate ammonification, and sulphate cycling. Genomic analysis of the most highly abundant members in this system suggests that an important source of chemotrophic energy is generated via the metabolic coupling of nitrogen and sulphur cycling.

**Conclusion:** These findings substantially contribute to our understanding of the novel and specialised microbial communities in anchialine ecosystems, and highlight key chemosynthetic pathways that appear to be important in these energy-limited environments. Such knowledge is essential for the conservation of anchialine ecosystems, and sheds light on adaptive processes in extreme environments.

## Introduction

The microbial communities of stratified aquatic systems serve as useful models for studying the relationships between metabolic strategies, water column depth, and physicochemistry. Stratified water columns, characterised by physical and chemical gradients, provide distinct niches for diverse assemblages of microbes, which, in turn, can support complex food webs in relatively extreme environments. Thus, unravelling the network of microbial metabolic strategies that link biogeochemical processes and trophic webs is important for understanding ecosystem functioning as well as evaluating ecosystem vulnerability [1].

Subterranean estuaries are stratified aquatic systems in which marine-derived groundwater mixes with meteoric freshwater in coastal aquifers [2]. These systems are globally distributed, and most commonly form in the porous limestone of karst coastlines [3]. They are characterised by water columns that exhibit stratified physicochemical profiles and low dissolved oxygen content [4]. Although they represent low-energy and extreme environments, subterranean estuaries can support complex ecosystems, which have been termed ‘anchialine’ [4]. The higher trophic levels of anchialine ecosystems largely comprise cave-adapted invertebrates with high rates of endemism [5, 6]. Earlier investigations into these anchialine food webs indicated that they may be supported, at least in part, by chemosynthetic microbes [7–9]. There is now growing interest in surveying the microbial communities that inhabit subterranean estuaries, and in particular, characterising their niche-adaptive metabolisms [1, 10]. Such endeavours are critical for assessing the vulnerability of anchialine ecosystems to environmental change.

Microbial ecology studies have revealed that anchialine ecosystems harbour highly diverse microbial assemblages. Examination of the prokaryotic community structure using 16S rRNA gene amplicon sequencing has been undertaken for several anchialine systems, including those found in Eastern Adriatic Sea Islands [11], Sansha Yongle Blue Hole in the South China Sea [12], Indonesian anchialine lakes [13], Blackwood Sinkhole in the Bahamas [14], and coastal aquifers of the Yucatán Peninsula, Mexico [10, 15]. These sites all revealed a high degree of taxonomic richness spanning functionally diverse microbial groups. Brankovits D*, et al.* [10] combined 16S rRNA gene sequencing with respiratory quinone biomarker analysis to infer the metabolic phenotypes of an anchialine water column, which contained a mixture of methanotrophs, heterotrophs, photoautotrophs, and nitrogen and sulphur cycling chemolithotrophs. They identified methane and dissolved organic carbon as key microbial energy sources that support higher trophic levels of the anchialine food web. Though, comparison between the microbial communities within coastal and in-land sinkholes of the same region (Yucatán Peninsula) show that the dominant metabolic strategies can differ significantly between different sinkholes along the same aquifer network [15].

Bundera Sinkhole, located in the karstic coast of Cape Range Peninsula in north-western Australia, is the only known continental anchialine system in the Southern Hemisphere. The sinkhole, which is the only opening to the subterranean estuary, is located 1.7 km inland from the Indian Ocean. The water column exhibits strong vertical stratification in its physicochemical profile, with decreasing dissolved oxygen and increasing salinity with depth, and polymodal peaks of inorganic nitrogen and sulphur compounds [16–18]. A range of endemic eukaryotes have been discovered in Bundera Sinkhole, including copepods, remipeds, and polychaetes [19–22]. Chemical profiling suggests that this trophic web may be supported by microbial chemosynthesis [16].

Microbial studies of Bundera Sinkhole using flow cytometry and 16S rRNA gene sequencing have shown the microbial communities to be stratified along the depth profile [17, 18, 23]. A diverse range of prokaryotes have been identified in the water column, comprising 67 identifiable bacterial and archaeal phyla [18]. Although community profiling suggests that a range of chemolithotrophic metabolisms are present throughout the water column, the high level of taxonomic novelty has made it difficult to infer the metabolic functions of many of the most abundant members [18]. Here, we employed shotgun metagenomic sequencing across a depth profile in Bundera Sinkhole to elucidate the metabolisms of these novel microbial communities. We identified key depth-dependent chemotrophic metabolic pathways, including coupled nitrogen-sulphur cycling, that may be driving nutrient feedback loops in this system. To the best of our knowledge, this is the first whole metagenomic sequencing approach of any anchialine ecosystem, and represents important findings that can help us to better understand microbial metabolic and biogeochemical processes in these unique environments.

## Methods

### Sample collection, DNA extraction and sequencing

Water samples were collected from Bundera Sinkhole as previously described [18]. Briefly, this involved pumping water samples from depths of 2, 8, 17, 18, 22, and 28 m between the 29th of June and the 1st of July 2015 for metagenomic analysis. For depths of 8 m and below, samples were collected using four previously installed boreholes (Fig. 1b). Physicochemical data, including salinity, dissolved oxygen (DO), dissolved organic carbon (DOC), ammonia (NH_3_), nitrate (NO_3_^−^), and sulphate (SO_4_^2−^) measurements were obtained from our previous study [18]. For metagenomic analysis, ∼4 L water samples were pre-filtered using 60 µm filters (Millipore Type NY60), and then passed through 0.2 μm Sterivex^TM^ filters. The 0.2 μm filters with captured microbial cells were cut from their casing, and DNA extractions carried out using the PowerWater® DNA Isolation kit (MO BIO Laboratories, Inc., Carlsbad, USA), according to the manufacturer’s protocol. Metagenomic libraries were prepared for duplicate biological replicates from each depth using the Illumina TruSeq DNA Library Preparation Kit, according to the manufacturer’s protocol, and sequenced on the Illumina HiSeq 2000 platform (High-Output v4).

**Fig. 1.**
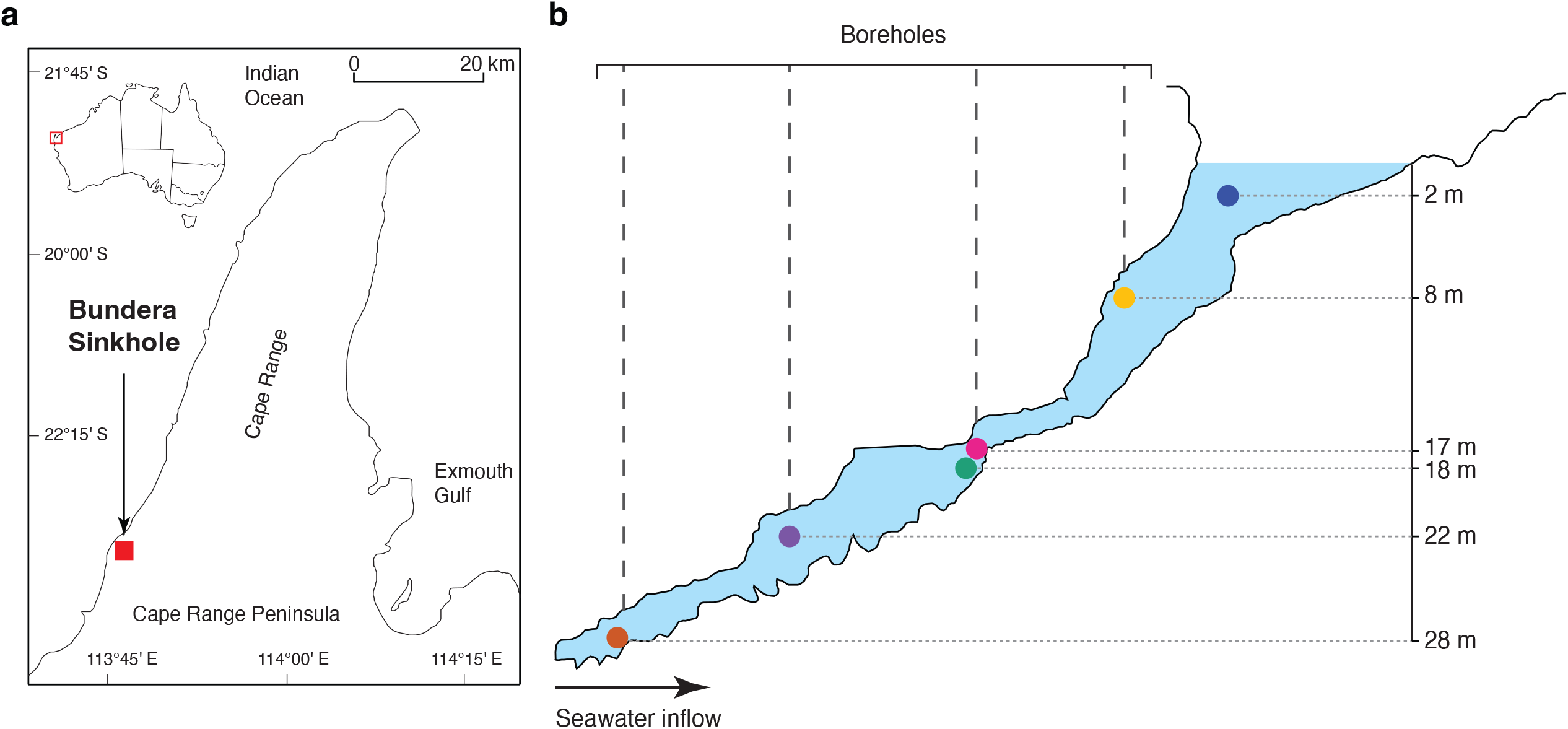
Location and sampling map of the Bundera Sinkhole. **(a)** Location of the Bundera Sinkhole in the Cape Range Peninsula, Western Australia. (**b**) Topology of the sinkhole and sampling points for shotgun metagenomic sequencing. Figure panels are adapted from Elbourne LDH*, et al.* [18].

### Metagenomic assembly and functional annotation

Raw reads were trimmed and quality filtered using Trimmomatic v 0.38 [24], and assembled with metaSPAdes v 3.13.0 [25] with default parameters. Quality of the assembly for each sample was assessed with QUAST v 5.0.2 using the metaQUAST option [26], and contigs shorter than 1 kb were removed from the assemblies. Open reading frames (ORFs) and translated protein sequences were predicted using Prodigal v2.6.3 [27] in metagenomic mode [parameter: -p meta]. ORFs from all samples were pooled and dereplicated at 98% nucleotide identity using CD-HIT v4.8.1 [28, 29] [parameters: -c 0.98 -n 10 -d 0 -t 0 -M 0]. The relative abundance of ORFs in each sample was calculated using the transcripts per million (TPM) method with CoverM v0.6.1 (https://github.com/wwood/CoverM) in contig mode [parameters: contig -t 24 --coupled -m TPM].

Translated protein sequences of the dereplicated ORFs were functionally annotated using METABOLIC v4.0 [30], by implementing the METABOLIC-G workflow with default parameters. The METABOLIC software identifies metabolic and biogeochemical traits by integrating several hidden Markov model (HMM) databases, comprising KOfam [31] (containing KEGG HMMs [32]), TIGRfam [33], Pfam [34], and custom [35] HMM databases.

### MAG binning and quality control

To improve sequencing depth, the replicate metagenome samples were co-assembled using MEGAHIT v1.2.9 [36, 37], with contig coverage calculated using Bowtie 2 v2.3.2 [38]. Co-assembled contigs were then binned using METABAT 2 v2.2.15 [39] with default parameters within Anvi’o v6.2 [40]. The resulting MAGs were then manually refined in Anvi’o. The completion and contamination of MAGs were estimated with CheckM v1.2.1 [41] using lineage-specific marker sets [parameters: lineage_wf -t 24]. MAG chimerism was assessed using GUNC v1.0.5 [42] with default parameters. Only MAGs that passed the GUNC chimerism check, had an estimated completion greater than 50%, and had an estimated contamination less than 10% were retained for further analysis. These represent the completion and contamination MIMAG criteria for high- and medium-quality MAGs [43].

### MAGs taxonomy and functional annotation

MAG taxonomy was assigned using GTDB-Tk v2.1.1 [44, 45] [parameters: classify_wf -- cpus 24] with release R207_v2 of the Genome Taxonomy Database (GTDB) [46–49]. We inferred domain-specific phylogenies using concatenated protein alignments generated by GTDB-Tk, which were based on the BAC120 [50] and AR53 [51] protein marker sets. The phylogenies were inferred from the alignments using a maximum-likelihood approximation employed by FastTree v2.1.10 [52, 53]. We applied a WAG substitution model with branch lengths rescaled to optimise the Gamma20 likelihood, and 1,000 resamples [parameters: - gamma -wag]. The inferred phylogenies were visualised using the ggtree v2.4.2 [54] and ggtreeExtra v1.7.0.990 [55] R packages.

MAGs were functionally annotated using METABOLIC v4.0 [30], by implementing the METABOLIC-C workflow with default parameters. The relative abundance of MAGs in each sample was calculated using the TPM method with CoverM v0.6.1 (https://github.com/wwood/CoverM) in genome mode [parameters: genome -t 24 --coupled - m TPM]. Four MAGs that were highly abundant, having TPM values greater than 50 in at least one sample, were further profiled for nitrogen cycling genes using the NCycDB [56]. DIAMOND v2.0.15 [57] was used to query MAG proteins against the NCycDB with a minimum E-value of 1e-05 [parameters: blastp -p 8 -k 1 -e 1e-5], and filtered using an amino acid identity cut-off of 70%.

### Statistical analyses

Beta-diversity analyses of the whole metagenomes, key metabolic genes, and MAG phyla were assessed using non-metric multidimensional scaling (NMDS) based on Bray-Curtis distances using the *vegdist* and *metaMDS* functions from the vegan v2.5-7 R package [58]. Groupings inferred from the NMDS ordination were compared with PERMANOVA using the pairwiseAdonis v0.4 R package [59], which uses the vegan functions, *vegdist* and *adonis*, to calculate inter-group differences in a pairwise fashion.

### Results and Discussion

Bundera Sinkhole, Australia’s only deep water anchialine system, supports a complex trophic web with an abundance of endemic micro- and macroorganisms. Previous chemical and community profiling using 16S rRNA gene sequencing suggest that this ecosystem may be sustained by microbial chemosynthesis [18, 23]. However, the high degree of taxonomic novelty, with associated uncertainty of metabolic functions, has limited our understanding of the dominant metabolic pathways in this system. Here, we employed shotgun metagenomic sequencing to investigate the distribution of key metabolic genes and to identify the biogeochemical cycling potential of the stratified microbial communities in Bundera Sinkhole.

### Microbial metabolic profiles are associated with water depth and physicochemistry

Bundera sinkhole exhibited a highly stratified water column with a marked physicochemical profile (Supplementary Table 1). The only oxic depth sampled was at 2 m, which had a dissolved oxygen (DO) concentration of 2.75 mg/L, and had the lowest salt concentration, with a salinity of 18.69 practical salinity units (PSS). The 8 m depth, representing the sinkhole’s halocline [16, 17], had a DO (0.86 mg/L) relatively higher than the samples from 17-28 m depths, and an intermediate salinity of 25.46 PSS. The lower depths, encompassing the 17-28 m samples, had lower levels of DO (0.28-0.47 mg/L) and higher salinity (31.41-32.35 PSS). Polymodal peaks of dissolved organic carbon (DOC), ammonia, nitrate and sulphate were observed along the water column (Supplementary Table 1).

Clear distinctions in microbial metabolic strategies were observed at different depths (Fig. 2). Microbial communities sampled from the 17, 18, 22, and 28 m depths exhibited similar metabolic gene diversity profiles, which differed from the 2 m and 8 m communities (Fig. 2b; PERMANOVA, P=0.04). The 2 m and 8 m metabolic profiles form distinct clusters based on NMDS analysis (Fig. 2b), although this separation was not determined to be significantly different (PERMANOVA, p=0.33), likely due to the limited statistical power of this comparison. The same clustering is observed for the beta-diversity of all genes detected in the metagenomes (Fig. 2a). Since genes were de-replicated at 98% nucleotide identity, clustering of all genes is more likely to reflect the taxonomic composition of the samples. Thus, both taxonomic and functional composition of the sinkhole appear to cluster according to salinity and oxygen concentrations. These same depth clusters are observed from16S rDNA amplicon sequencing of the sinkhole [18].

**Fig. 2.**
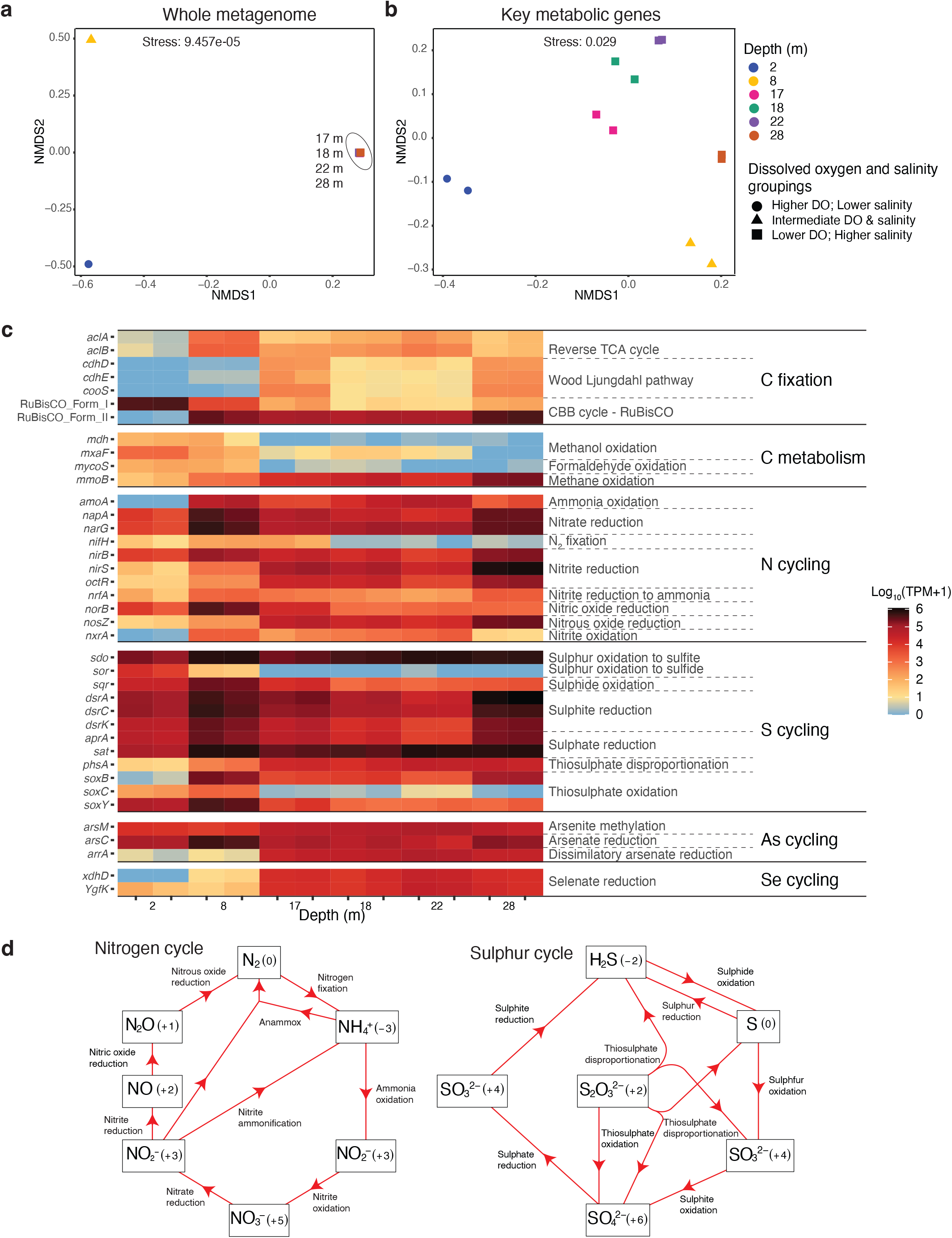
Relative abundance and diversity of key metabolic and biogeochemical cycling genes in Bundera Sinkhole. **(a-b)** Non-linear multidimensional scaling (NMDS) based on Bray-Curtis distances of normalised read counts for (**a**) whole metagenomes (with genes dereplicated at 98% nucleotide identity) and (**b**) key metabolic genes (TPM sums) displayed in panel **c**. In panel **a**, NMDS points that represent replicate samples lie on top of each other, as do those representing all samples from 17, 18, 22, and 28 m depths. The NMDS groupings (circles, triangles, and squares) represent samples with similar levels of dissolved oxygen (DO) and salinity (Supplementary Table 1). In both NDMS plots, the grouping of samples from 17, 18, 22, and 28 m depths (squares) is supported by PERMANOVA (p=0.04; Supplementary Table 2). (**c**) Relative abundance of key metabolic marker genes within each sample. Colour scale is displayed as log_10_(TPM +1) to account for TPM values of zero. Gene names are displayed to the left of the heatmap, and the reactions that they facilitate are on the right. (**d**) Visualisation of microbial nitrogen and sulphur cycling pathways present in Bundera Sinkhole. Chemical compounds that represent either the substrate or product of a reaction are boxed, with oxidation states shown in parentheses.

Autotrophic CO_2_ fixation strategies differed by depth (Fig. 2c), likely in response to oxygen levels and percentage of incident light. The CBB cycle, which utilises the CO_2_fixation enzyme ribulose-1,5-bisphosphate carboxylase/oxygenase (RuBisCO) by photo- and chemo-autotrophs, was depth-dependent. Two main forms of RuBisCO are known to be involved in the classical CBB cycle [60]. Surface samples, particularly those from the 2 m depth, were characterised by a greater relative abundance of the Form I RuBisCO compared to other depths (and other C-fixation strategies), presumably from a greater abundance of photoautotrophs. While the relative abundance of form II RuBisCO, which is adapted to low-O_2_ conditions [61], had an opposite trend, with greater relative abundance at lower depths. The relative abundance of genes that drive the reverse TCA cycle and Wood-Ljungdahl pathway increased with depth, which are the hypoxic regions of this system (Supplementary Table 1). Similar trends have been observed in hypoxic and anoxic zones of stratified water columns [62, 63].

The relative abundance of marker genes for different pathways involved in carbon metabolism also corresponded to a depth gradient (Fig. 2c). Methanol and formaldehyde oxidation (C1 metabolism), decreased with depth, while the methane monooxygenase gene, *mmoB*, involved in the first step of methane metabolism, increased with depth. Similar patterns of carbon metabolism genes have been observed over an oxygen gradient in a permanently stratified lake [63]. Arsenic and selenium cycling genes also corresponded to a depth gradient (Fig. 2c). In particular, the abundance of genes involved in dissimilatory (respiratory) arsenate and selenate reduction increased with depth. Both arsenate and selenate can be utilised in anerobic respiration for energy production [64, 65], explaining their greater relative abundance at hypoxic depths. These elements can thus provide additional energy sources for facultative or obligate anaerobes at the lower depths of the sinkhole.

Pathways for the complete cycling of nitrogen (N) and sulphur (S) compounds were observed in the sinkhole (Fig. 2d), with diverse N and S cycling reactions present at different depths (Fig. 2c). Several key N and S cycling genes were strongly correlated with concentrations of ammonia, nitrate, and sulphate (Fig. 3; Supplementary Table 3), highlighting these as key environmental parameters. To infer the direction of these correlations and to identify nutrient feedback loops, we examined whether the correlated genes were involved in either the production or substrate utilisation of these chemical compounds. Marker genes for N cycling that correlated with ammonia concentrations were all involved in pathways that produced ammonia (Fig. 3a-c; Supplementary Table 3). These included: *napA*, encoding a nitrate reductase, involved in the first step of the dissimilatory nitrate reduction to ammonia (DNRA) pathway, reducing nitrate to nitrite; and *nirB* and *nrfA*, both encoding nitrite reductases, involved in the second step of the DNRA pathway, further reducing nitrite to ammonia. Similarly, N cycling marker genes that correlated with nitrate concentrations were all involved in nitrate production (Fig. 3d,e; Supplementary Table 3). These included *amoA*, encoding an ammonia monooxygenase, involved in the first step of nitrification, oxidising ammonia to nitrite; and *nxrA*, encoding a nitrite oxidoreductase, involved in the final step of nitrification, oxidising nitrite to nitrate. We also found that the relative abundance of both *amoA* and *nxrA* are negatively correlated with the concentration of dissolved organic carbon (DOC) (Fig. S1; Supplementary Table 4), suggesting that chemolithotrophic nitrification is an important metabolic pathway when available organic carbon is limited. Thus, microbial communities and environmental concentrations of DOC, ammonia and nitrate are apparently linked in a feedback loop involving nitrification (ammonia to nitrate) and DNRA (nitrate to ammonia) pathways.

**Fig. 3.**
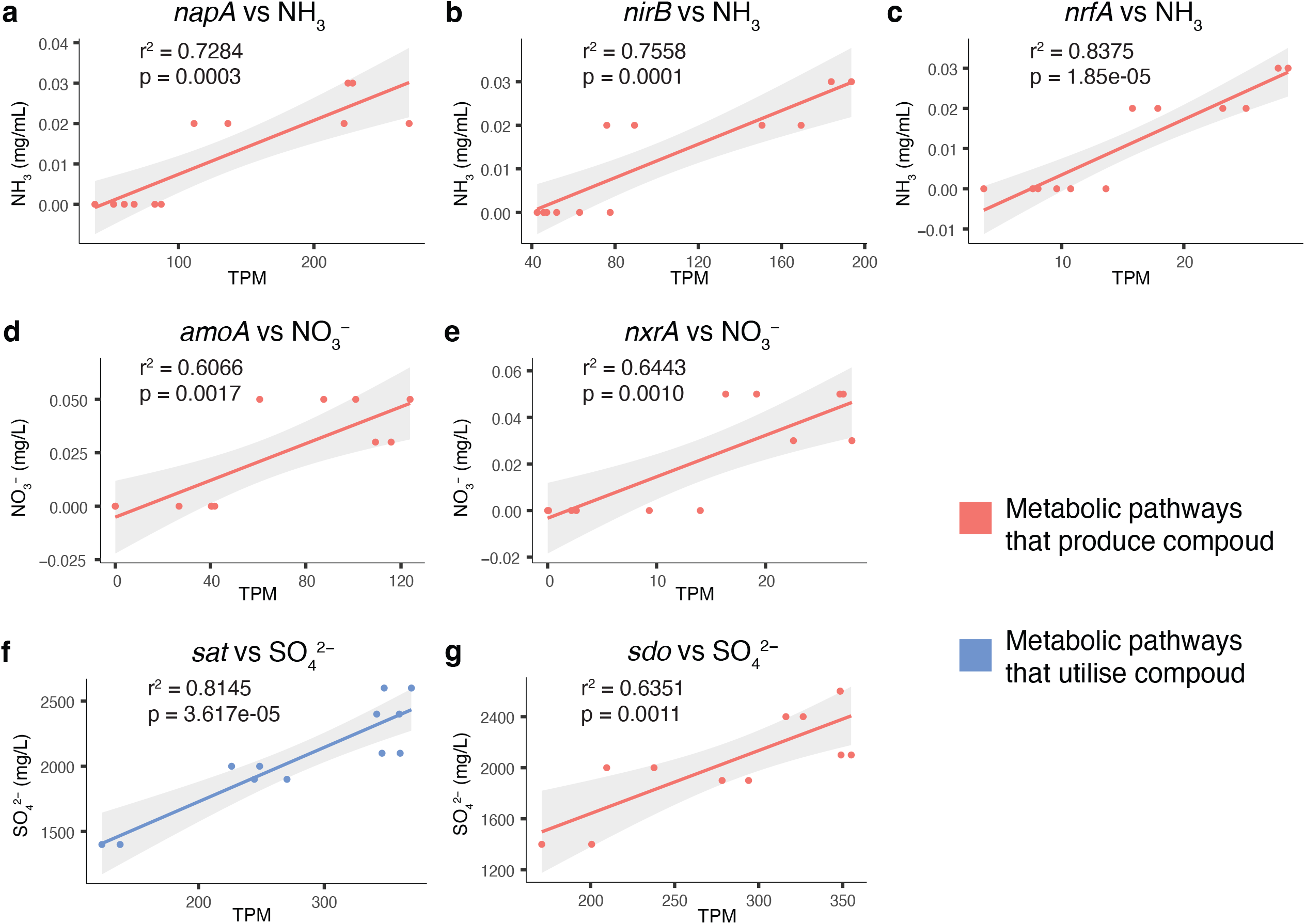
Correlations between chemical compound concentrations and genes involved in their cycling. Nitrogen and sulphur cycling genes whose relative abundance (TPM) are strongly correlated (r^2^ > 0.6) with the environmental concentrations of (**a**-**c**) ammonia (NH_3_), (**d**-**e**) nitrate (NO ^−^), and (**f**-**g**) sulphate (SO ^2−^). Plots coloured red represent genes involved in pathways that produce the corresponding chemical compound, either directly (**b**,**c**,**e**) or indirectly, via an intermediate compound (**a**,**d**,**g**). Correlation between *sat* gene relative abundance and SO4 concentrations (**f**) is coloured blue to indicate the gene’s involvement in SO4 substrate utilisation. Shaded regions represent the 95% confidence interval of the fitted linear model. A full list of r^2^ and p-values for all evaluated nitrogen and sulphur cycling gene correlations is presented as Supplementary Table 3.

The S cycling marker genes, *sat* and *sdo*, were significantly correlated with sulphate concentrations (Fig. 3f,g; Supplementary Table 3), and are involved in the utilisation and production of sulphate, respectively. *sat* encodes a sulphate adenylyltransferase that coverts sulphate to adenosine-5′-phosphosulfate (APS) [66]. *sdo* encodes a sulphur deoxygenase which oxidises glutathione persulphide (GSSH). Sulphite is the first product of SDO activity via GSSH oxidation, which then leads to the non-enzymatic production of sulphate (likely from auto-oxidation of sulphite) [67]. Thus, sulphate concentrations in Bundera Sinkhole are likely being driven by, as well as shaping, the microbial communities in a sulphate-feedback loop.

### Taxonomically novel and functionally diverse prokaryotes inhabit the sinkhole

Bundera Sinkhole harbours considerable microbial diversity, so to gain better insight into the metabolic potential of the novel and abundant microbial species, we employed genome-resolved metagenomic analysis. We generated 180 medium- to high-quality MAGs from the twelve co-assembled metagenomes (median completion = 88.75%, median contamination = 0.93%; Supplementary Table 5). These comprised 150 bacterial MAGs from 20 phyla, with the remaining 30 MAGs from 3 archaeal phyla (Fig. 4). The composition of prokaryotic phyla differed significantly by water depth, with distinct phyla found at 2 m, 8 m, and 17-28 m depths (Fig. S2; Supplementary Table 6), reflecting the same groupings as the gene-based clusters. This is supported by 16S rDNA amplicon sequencing of Bundera Sinkhole communities [18], which suggests similar depth-dependent composition of microbial taxa.

**Fig. 4.**
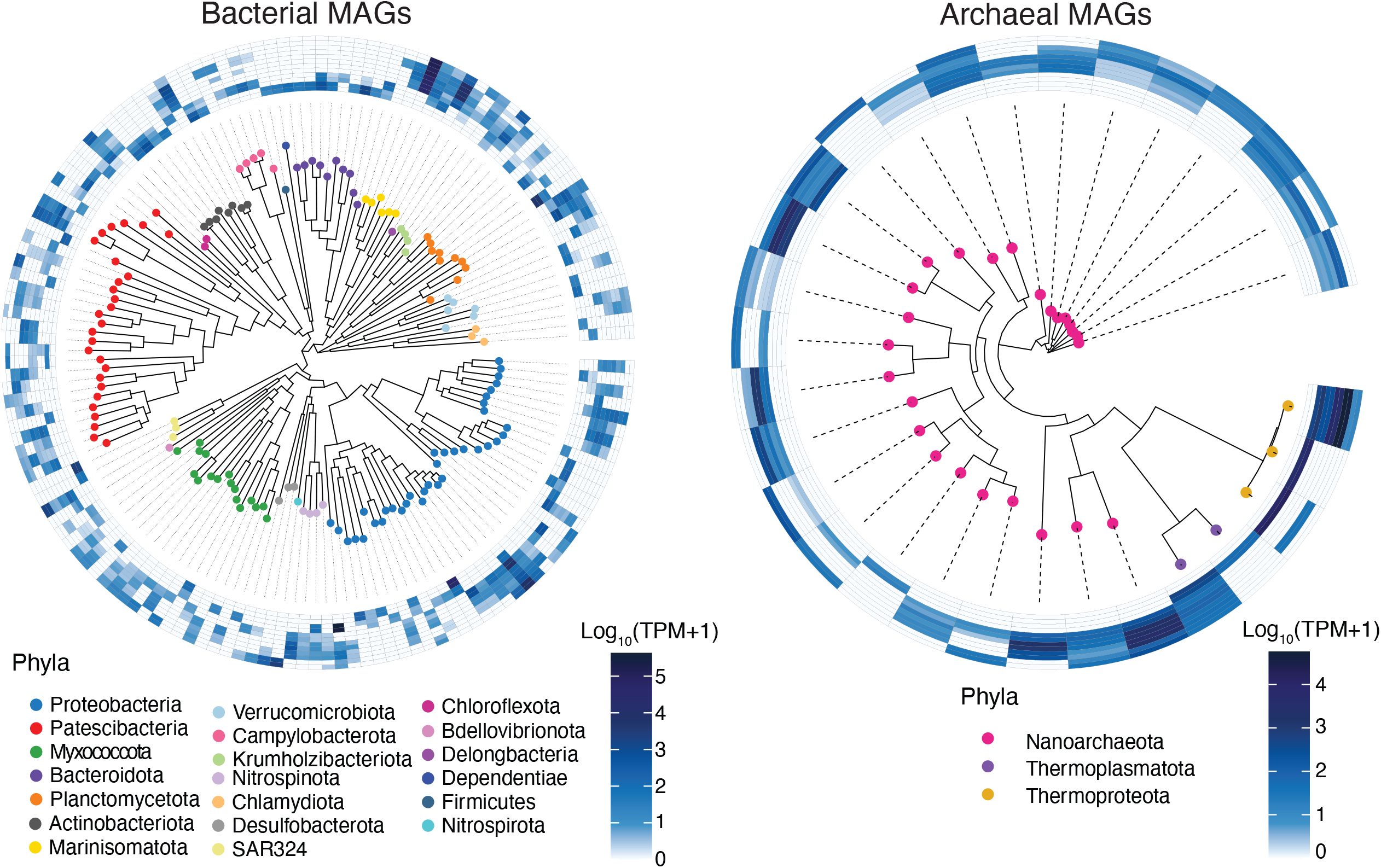
Domain-specific phylogenies of MAGs from Bundera Sinkhole. Tips of the trees are coloured by their assigned phylum. Heatmaps display the relative abundance of MAGs in each of the duplicate samples collected from six depths (from inner to outer rings: 2 m, 8 m, 17 m, 18 m, 22 m, and 28 m).

The communities inhabiting Bundera Sinkhole are taxonomically novel, with 75% of MAGs assigned to entirely new or uncharacterised families that lack cultured representatives. In the Genome taxonomy Database (GTDB), newly delineated taxa are allocated with alphanumeric placeholder labels. Using GTDB nomenclature, we found that 64% of MAGs were assigned to families with such placeholder labels, and a further 11% of MAGs could not be assigned to any family (Supplementary Table 5). Even at the class level, almost a quarter of all MAGs in this system were assigned to placeholder-labelled lineages. Such taxonomic novelty is likely driven by niche adaptation to the distinctive geomorphological and physicochemical properties of anchialine ecosystems.

The suite of MAGs assembled from Bundera Sinkhole provides an ideal opportunity to assess the functional potential of these diverse and novel taxa. The relative abundance of MAG-related functions associates with water depth (Fig. 5), as observed with the gene-based functional analysis. We found that the number of MAGs that have the genetic potential for each key metabolic reaction varied considerably, as does their relative abundance at different depths.

**Fig. 5.**
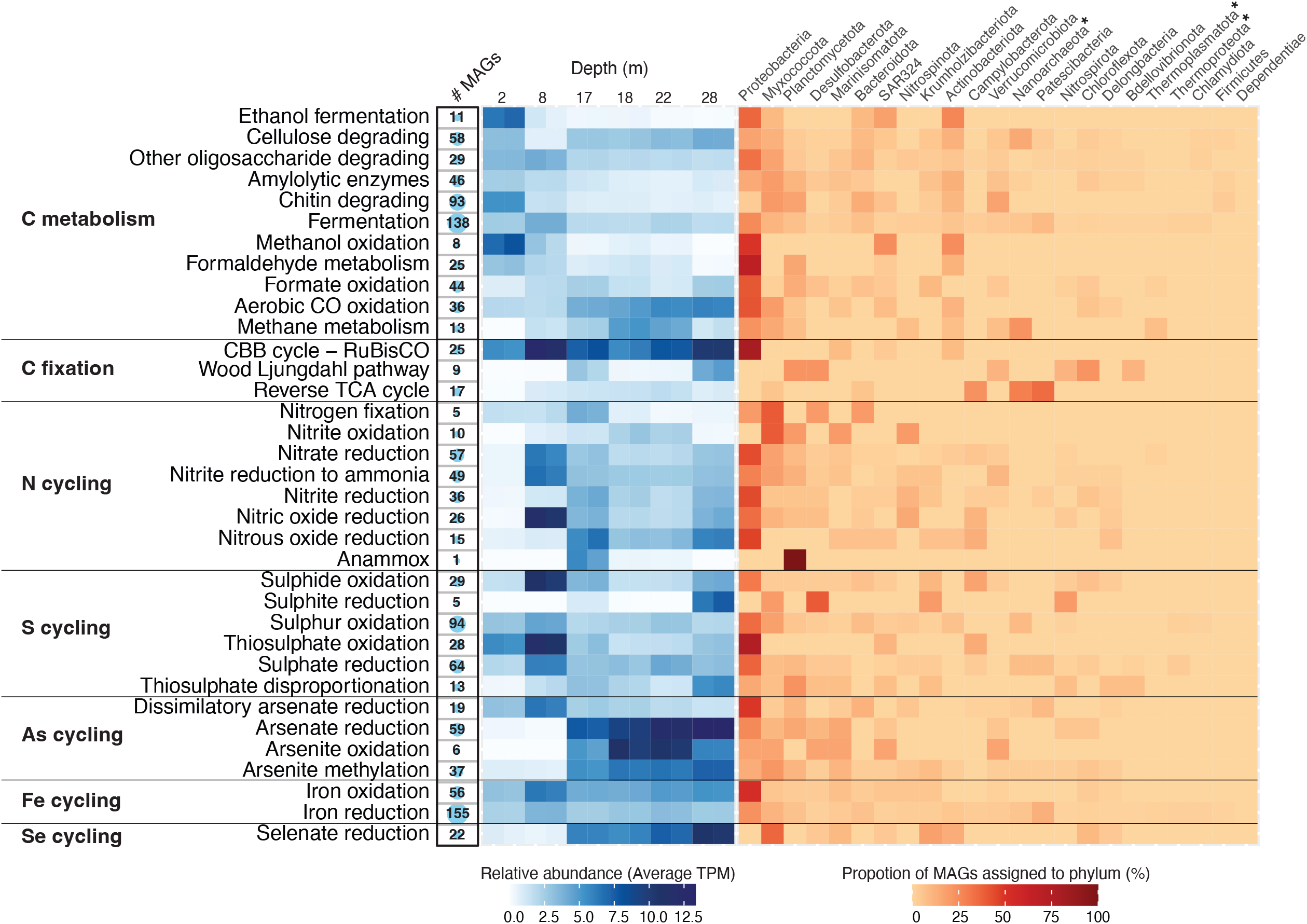
Key metabolic and biogeochemical cycling traits of MAGs in Bundera Sinkhole. From left to right: the numbers of MAGs that carry genetic markers (listed in Supplementary Table 7) for each functional trait are displayed by numerals, and represented visually by the size of the circles; the average relative abundance (TPM) for corresponding MAGs at each depth are displayed by the blue heatmap; and the proportion of MAGs assigned to each phylum is represented by the red heatmap. Archaeal phyla are denoted with asterisks.

We found that the taxonomy of carbon metabolism varied based on the carbon substrate (Fig. 5). For example, one-carbon (C1) molecules (e.g., methanol, formaldehyde, formate, and carbon monoxide) are largely metabolised by Proteobacteria, while complex carbon molecules (e.g., cellulose, chitin, starch, and other oligo- and poly-saccharides) are metabolised by bacteria from a wider range of phyla.

The taxonomy of autotrophic microbes differed based on the CO_2_ fixation strategy (Fig. 5). Photo- and chemo-autotrophs that utilise RuBisCO as part of the carbon-fixing CBB cycle were almost all Proteobacteria (80%). A much more diverse range of bacteria and archaea had the genetic potential for utilising the reverse TCA (Patescibacteria, Nanoarchaeota, Campylobacterota, Myxococcota, Bacteroidota) and Wood-Ljungdahl (Planctomycetota, Desulfobacterota, Chloroflexota, Verrucomicrobiota, Nitrospirota, Bdellovibrionota) pathways for carbon fixation.

For the most part, N and S cycling pathways were performed by Proteobacteria (Fig. 5). As described above, both the DNRA and nitrification processes appear to be important N cycling pathways that drive a nitrogen-feedback loop in this system. The DNRA pathway, involving nitrate reduction to nitrite, which is then further reduced to ammonia, is largely driven by Proteobacteria (Fig. 5). The reverse of this process, nitrification, involves ammonia oxidation to nitrite, which is further oxidised to nitrate. Here, the final nitrification step (nitrite oxidation) is predominately driven by Myxococcota, and to a lesser extent, Planctomycetota, Marinisomatota, and Nitrospinota (Fig. 5). However, the first step in nitrification (ammonia oxidation), mediated by ammonia monooxygenases, was not detected in any MAG, despite their presence in the gene-based analysis (Fig. 2c). Therefore, to identify the taxa involved in ammonia oxidation, we queried the genes annotated as *amoA* (encoding the ammonia monooxgenase, alpha subunit) against NCBI’s nr database using BLASTP. Three *amoA* genes were detected among the set of de-replicated genes. All three were identified as archaeal, belonging to the NCBI phylum Thaumarchaeota (classified in the GTDB as class Nitrososphaeria – phylum Thermoproteota [48]). Thus, the nitrogen-feedback loop that cycles between ammonia and nitrate is driven by distinct prokaryotes – predominately those belonging to Proteobacteria, Myxococcota, and Archaea. The aforementioned sulphate-feedback loop, associated with sulphate reduction (*sat*) and sulphur oxidation (*sdo*) processes, is also largely driven by Proteobacteria (Fig. 5).

Given the large metabolic contribution of Proteobacteria to this system, we further investigated their functional potential at lower taxonomic levels (Fig. 6). We found that the most important contributors to key metabolic reactions (based on read coverage) are species from less well characterised proteobacterial lineages. In particular, bacteria belonging to the gammaproteobacterial orders PS1 (n=1) and GCF-002020875 (n=7) were key contributors to carbon fixation (CBB cycle), and nitrogen and sulphur cycling (Fig. 6). The single PS1 MAG belongs to the genus *Thioglobus*, which encompass members of the sulphur-oxidising marine SUP05 clade of Gammaproteobacteria. *Thioglobus* comprises a handful of cultured representatives which consist of chemoauto- and hetero-trophic bacteria that grow under aerobic and anaerobic conditions, and are assumed to contribute to denitrification [68–71]. The seven MAGs assigned to the order GCF-002020875, which lacks any cultured representatives, all belong to the same family, also designated GCF-002020875. Of these, four MAGs belong to the genus *Thiopontia*, while the other three MAGs were unclassified at the genus level. There are five species representative MAGs for *Thiopontia* (GCA_018671205.1, GCA_018658305.1, GCA_018648825.1, GCA_013349825.1, GCA_014384675.1), all of which were assembled from hypoxic saline water metagenomes [72–74] (NCBI BioProject Accessions: PRJNA630981, PRJNA632036, and PRJNA649215), suggesting that these bacteria are specific to this environmental niche.

**Fig. 6.**
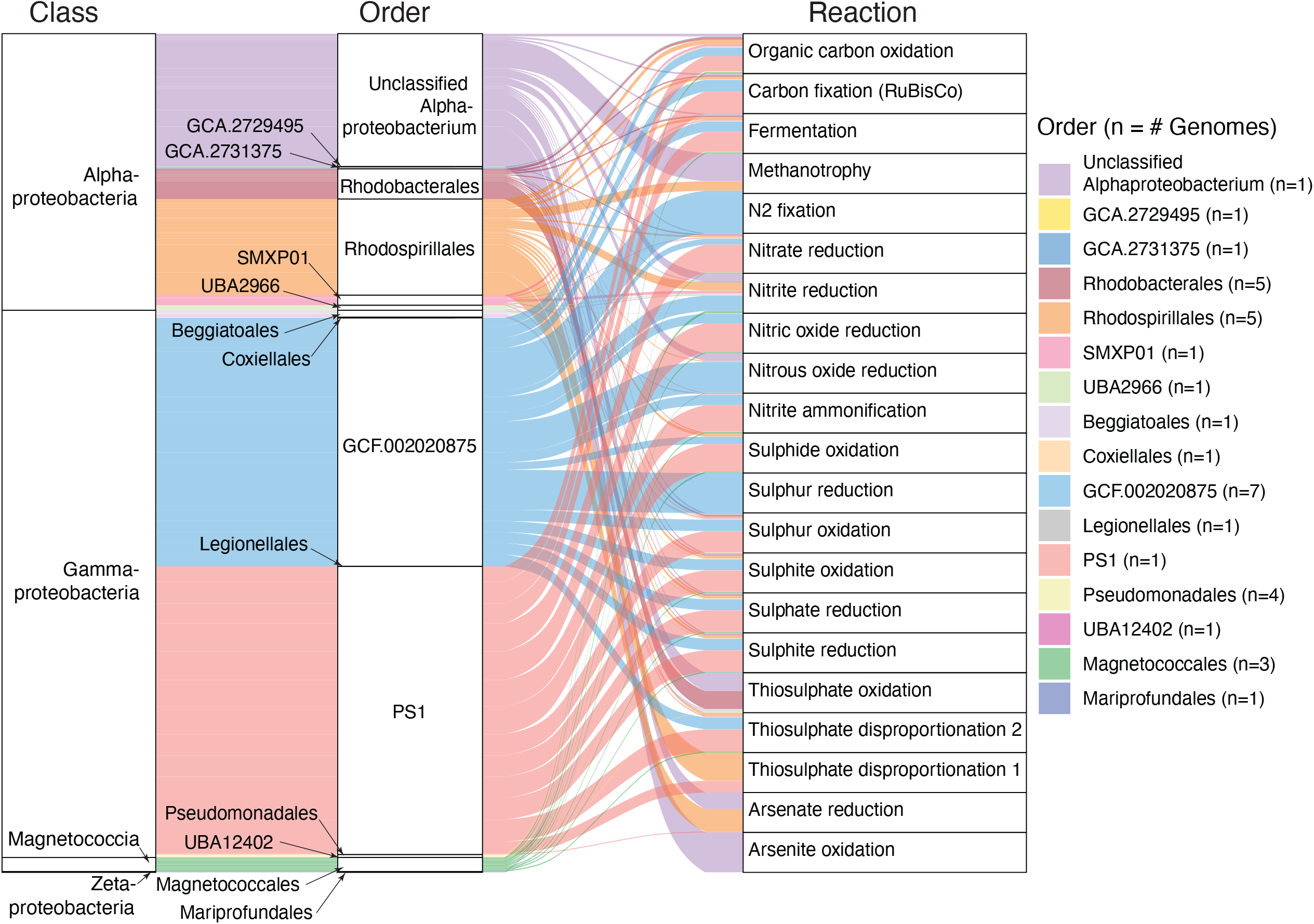
Metabolic functions associated with proteobacterial MAGs. MAGs are grouped according to their taxonomic class (left) and order (middle). Width of curved lines indicate the relative contribution, based on read coverage, of proteobacterial orders (middle) to a given metabolic reaction (right).

### Bundera Sinkhole has one to two highly abundant MAGs at each depth

Four highly abundant MAGs (with TPM values >50 in at least one sample) were dominant at different depths (Fig. 7). These included two gammaproteobacterial MAGs, one assigned at the family level (family GCF-002020875), and a *Thioglobus* sp., which were highly abundant at the 2 m and 8 m depths, respectively. A Marinisomatota MAG (order Marinisomatales) was highly abundant across all lower-depth samples (17-28 m). An archaeal MAG, *Nitrosopumilus* sp., was also abundant across the lower-depth samples, particularly, at the 22 m depth.

**Fig. 7.**
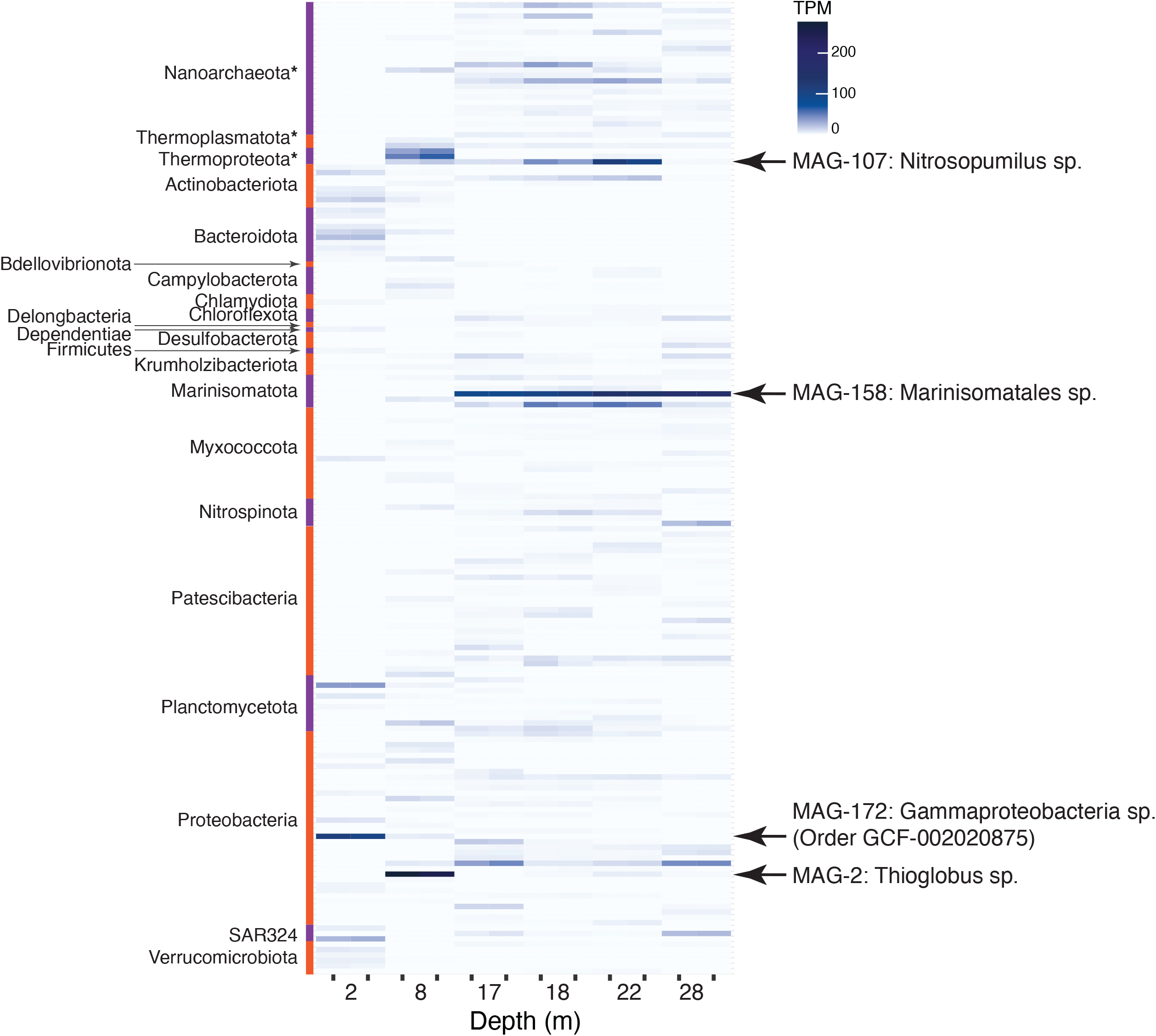
Relative abundance of MAGs in Bundera Sinkhole. Phyla of MAGs are displayed to the left of the heatmap. Archaeal phyla are denoted with asterisks. The four most abundant MAGs, having a TPM value greater than 50 in any one sample, are denoted on the right.

The GCF-002020875 MAG (MAG-172), which comprised ∼9% of the metagenomic reads from the 2 m samples (Fig. 8), represents a novel gammaproteobacterial lineage, having no classification below the family level. It encodes several enzymes that would enable it to utilise sulphur as an energy source. However, it also carries genes for complex carbon degradation, suggesting it has the potential for both thioauto- and hetero-trophy. It also has the genetic potential to mediate two steps in the denitrification pathway (nitrite reduction to nitric oxide, and nitrous oxide reduction to N_2_ gas).

**Fig. 8.**
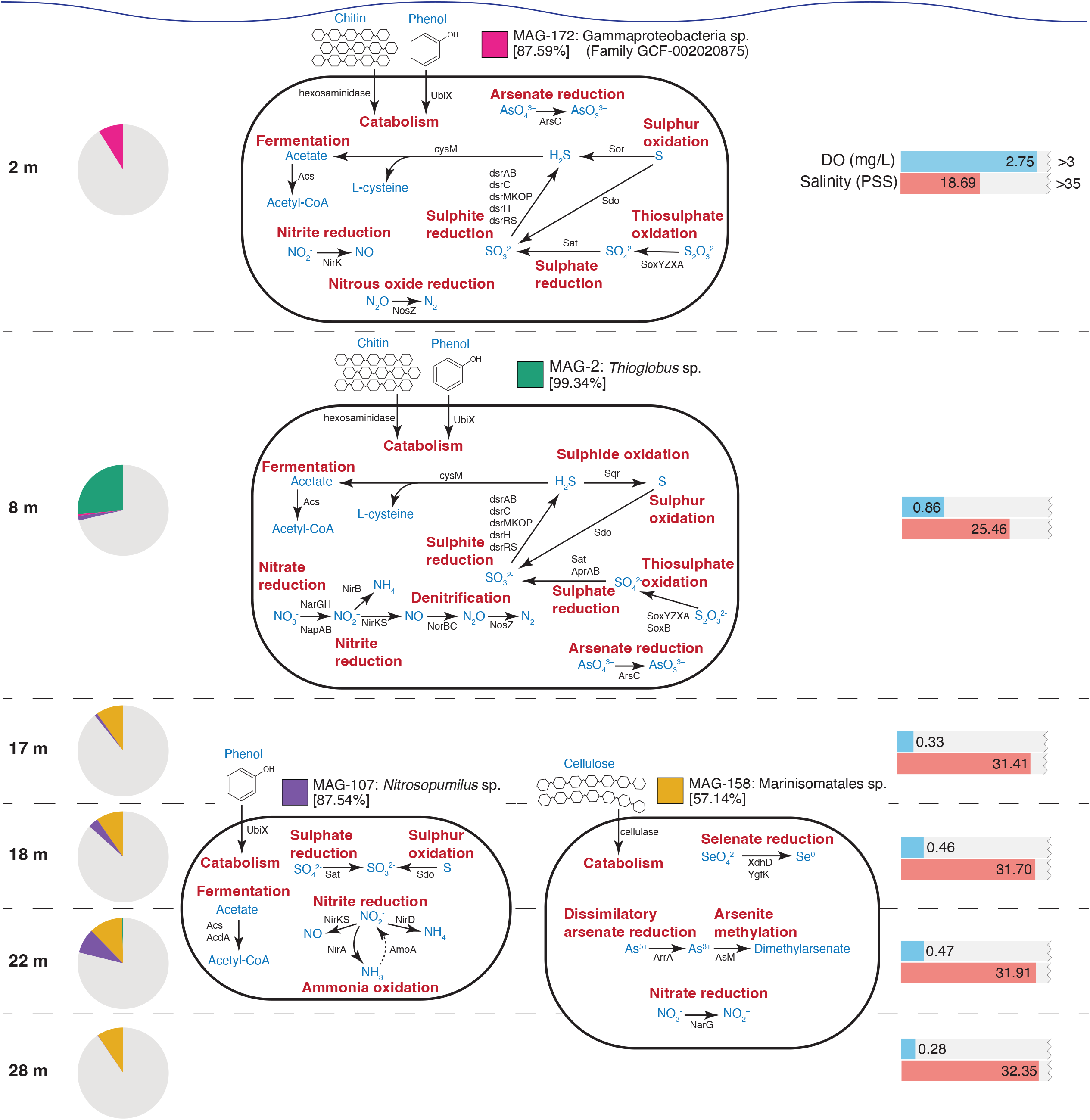
Metabolisms of the most highly abundant MAGs in Bundera Sinkhole. Estimated genome completeness is displayed within square brackets under each MAG ID. Pie charts indicate the proportion of reads at each depth that map to the four MAGs. Metabolic reactions are labelled in red text, proteins mediating those reactions are labelled in black text, and the reaction products/substrates are labelled in blue text. Bar charts indicate the dissolved oxygen (DO) and salinity at each depth. In MAG-107, ammonia oxidation is displayed as a dashed arrow, as the *amoA* gene was not originally binned with this MAG. However, it was included here after detecting an *amoA* gene, taxonomically classified as *Nitrosopumilus*, that had a relative abundance almost perfectly correlated (r^2^ = 0.97) with that of MAG-107.

The highly abundant *Thioglobus* MAG (MAG-2) represents a major component of the 8 m community, comprising 26% of the reads from the 8 m samples (Fig. 8). It encodes several enzymes that suggest it also has the capacity for both thioauto- and hetero-trophy. It appears to be an important mediator of sulphur cycling, encoding several sulphur transformation pathways, and carries marker genes for the complete denitrification pathway, converting nitrate to N_2_ gas, via nitrite, nitric oxide, and nitrous oxide intermediates. Both dominant MAGs at the 2 m and 8 m depths possess the genetic potential for several sulphur cycling pathways as well as denitrification (Fig. 8). In marine oxygen minimum zones, a denitrification pathway linking reduced sulphur compounds to the loss of bioavailable nitrogen represents an important mode of metabolic coupling [75–78]. These two dominant MAGs are likely mediating this linking of sulphur cycling and denitrification in the shallower waters of the sinkhole.

In the deeper layers (17-28 m), two MAGs were highly abundant. One of these, MAG-107, belongs to the genus *Nitrosopumilus*, which comprise a group of ammonia-oxidising Archaea [79]. Given their important ecological role in ammonia oxidation, we searched this MAG for the marker gene for ammonia oxidation, *amoA*, encoding the ammonia monooxygenase alpha subunit. Surprisingly, *amoA* was not detected in this MAG. However, as described above, we detected three archaeal *amoA* genes from the complete set of de-replicated metagenomic genes. One of these was predicted to belong to the genus *Nitrosopumilus* (100% query cover and 98.61% amino acid identity to *Nitrosopumilus* AmoA [NCBI accession WP_141977518.1]), and its relative abundance is almost perfectly correlated (r^2^ = 0.97) with that of MAG-107, suggesting it to be indeed a component of its genome. The failure for the *amoA* gene to be binned with MAG-107, is possibly due to the several ribosomal protein genes co-located on the same contig (*rpl32e*, *rpl19e*, *rpl10*, *rpl12*, *rpl21e*, *rps17e*, *rps11*, *rps15*, *rps3ae*), which are often difficult to bin because of their differential codon usage patterns that have been optimised for rapid translation [80]. Besides ammonia oxidation, this MAG also had the genetic potential for several nitrate reduction pathways, as well as sulphite production (Fig. 8).

The Marinisomatales MAG, MAG-158, represents the other dominant MAG at the lower depths. This MAG belongs to the phylum Marinisomatota, also commonly known as Marinimicrobia. These bacteria are widespread in the global oceans, and are particularly abundant in sub-euphotic oxygen minimum zones [75], which correspond to the samples that MAG-158 was most abundant. Out of the four dominant MAGs, MAG-158 had the lowest estimated genome completeness (57.14%), partially obscuring detailed analysis of its metabolism. Nevertheless, we detected several enzymes involved in selenium and arsenic cycling, as well as nitrate reduction (representing the first step in denitrification) (Fig. 8). Previous analyses of these bacteria indicate that they are important drivers of denitrification and sulphur cycling in hypoxic and anoxic seawater [75, 81], suggesting that this MAG might also be involved in coupled sulphur-nitrogen cycling in the sinkhole.

## Conclusion

Here, we characterised the metabolic and biogeochemical cycling potential of the microbial communities inhabiting Bundera Sinkhole. We found that the microbial communities, largely represented by novel taxonomic lineages, display depth-dependent metabolisms. Key metabolic genes group into three depth-specific clusters that reflect distinct phases along the dissolved oxygen and salinity gradients. In particular, chemotrophic metabolisms that couple nitrogen and sulphur cycling appear to be characteristic of the dominant members in this ecosystem. These data support the idea that microbial chemosynthesis is sustaining the higher trophic levels in the sinkhole. To the best of our knowledge, this is the first whole metagenomic analysis of an anchialine ecosystem, and thus presents key findings that contribute to our understanding of ecosystem functions in subterranean estuaries.

Understanding the diversity of metabolic strategies utilised by anchialine microbial communities can provide important insights into how trophic webs are supported in these unique ecosystems. This is particularly important given the high endemism of anchialine species and the potential vulnerability of these ecosystems to global environmental change and other anthropogenic influences [1]. Identifying the key microbial members and biogeochemical process is critical for the conservation of anchialine ecosystems.

## Ethics approval and consent to participate

Not applicable

## Consent for publication

Not applicable

## Availability of data and material

Raw metagenomic sequence data are available in the NCBI SRA Database under BioSample Accessions SAMN32209613-SAMN32209624, from the BioProject PRJNA911846.

## Authors’ contributions

T.M.G conducted the data analyses and wrote the manuscript draft. A.F and L.D.H.E conducted data analyses. B.S performed the experimental work. W.H collected the water samples and was involved in the project design. I.T.P and S.G.T were involved in project design and management. All authors contributed to the final editing of the manuscript.

## Acknowledgements

TMG would like to thank Mary Ghaly for support and discussions regarding the manuscript.

## Supplementary Figures

**Fig. S1.**
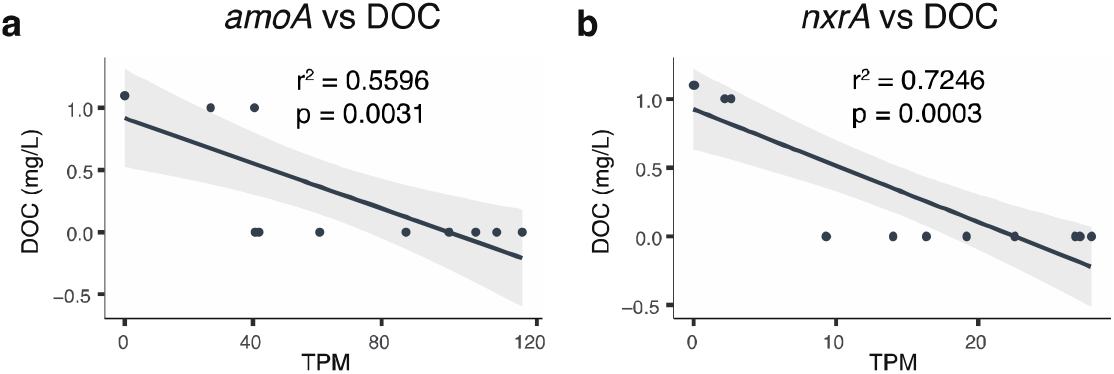
Correlation between dissolved organic carbon and nitrification. **(a)** Correlation between the relative abundance (TPM) of the marker gene for ammonia oxidation (*amoA*; first step of nitrification) and dissolved organic carbon (DOC) concentration. (**b**) Correlation between the relative abundance of the marker gene for nitrite oxidation (*nxrA*; final step of nitrification) and DOC concentration. Shaded regions represent the 95% confidence interval of the fitted linear model. A full list of r^2^ and p-values for all evaluated nitrogen and sulphur cycling gene correlations is presented as Supplementary Table 4.

**Fig. S2.**
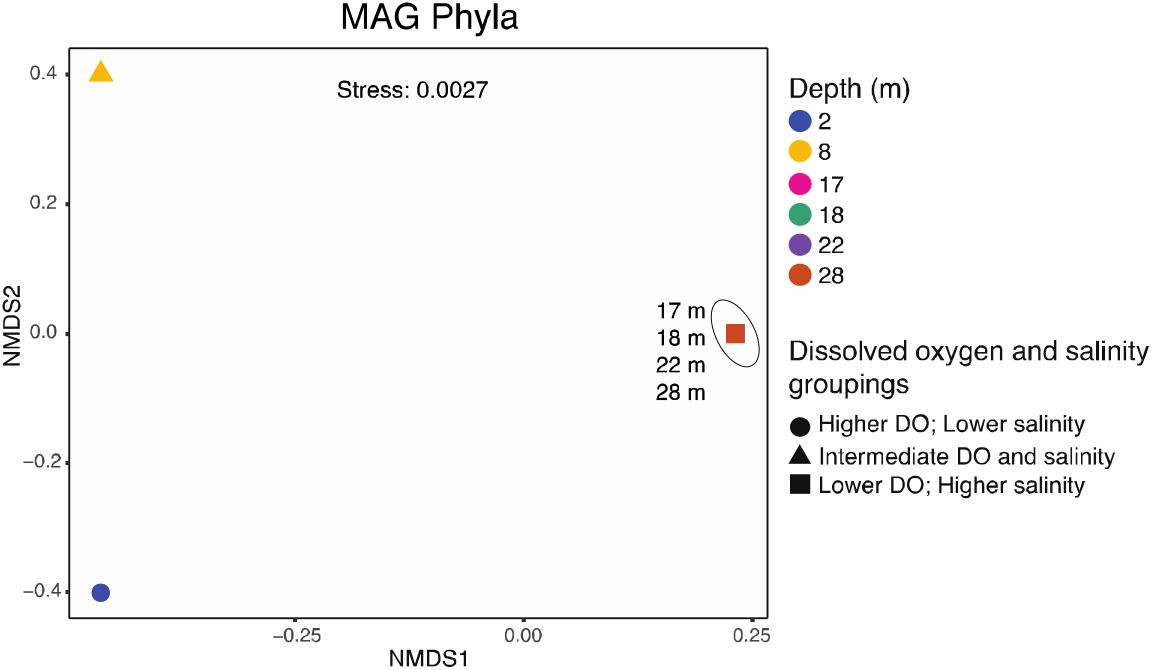
Beta-diversity of MAG phyla in the Bundera sinkhole. Non-linear multidimensional scaling (NMDS) based on Bray-Curtis distances of normalised read counts for MAG phyla. NMDS points that represent replicate samples lie on top of each other, as do those representing all samples from 17, 18, 22, and 28 m depths. The groupings (circles, triangles, and squares) represent samples with similar levels of dissolved oxygen (DO) and salinity (Supplementary Table 1). The grouping of samples from 17, 18, 22, and 28m depths (squares) is supported by PERMANOVA (p=0.046; Supplementary Table 6).

